# Developmental, time-of-day, and stimulus-specific Golgi-Cox staining patterns detected in the mouse brain

**DOI:** 10.1101/2025.07.03.663022

**Authors:** Melissa E. S. Richardson, Lauren Michel Barnwell

## Abstract

Well-coordinated brain activity is a crucial driver of bodily functions and is refined by environmental input. Understanding the structure of brain areas that regulate various functions and how the environment affects the neural responses in the brain has been fundamental in advancing the field of neuroscience and medicine. The Golgi-Cox method is a histological approach that has allowed researchers to view neuronal structures with unmatched and detailed resolution, allowing for the comparison between non-diseased and diseased models, for instance. However, this method is known to stain neurons sparsely, which is useful for distinguishing structural components, but unpredictably, which is difficult for reproducibility and targeted studies. Here, we use three approaches to demonstrate a predictable pattern of cell staining using the Golgi-Cox method. We show that neuronal maturity, time of day, and response to environmental stimuli affect the number of cells stained by the Golgi-Cox method. Furthermore, we found low variability within each experimental group, which indicates staining reproducibility under controlled environments. Our study highlights important parameters for using the Golgi-Cox method and demonstrates its feasibility for broader application in answering neuroscience-based questions.

**Summary:** Our study provides previously unknown insights demonstrating that the historical Golgi-Cox staining pattern of neurons is specifically linked to age, time-of-day, and light-responsiveness at night in the mouse brain.

## Introduction

Over the years, numerous methods have been devised to study the brain’s molecular, structural, and physiological changes during development and in response to environmental changes. Some methods include C-fos immunohistochemistry (IHC)(to assess neuronal activity) (Lara Aparicio *et al*., 2022), BrdU IHC (to determine the presence of newborn neuronal cells) (Yu *et al*., 1992), antigen-specific antibodies (to identify protein expression in specific cell types) (De Matos *et al*., 2010), Nissl staining (to identify neuron cell bodies), and Golgi staining (to identify neuronal structure) (Ghosh and Walocha, 2024). These methods vary with the specificity of the types of neurons they stain and cost-effectiveness (Flores *et al*., 2015; Zhong *et al*., 2019; Golberg *et al*., 2024).

The Nissl and Golgi staining are widely recognized as methods that are both cost-effective and able to capture location and morphological structures (respectively) of neurons in multiple brain regions (Einarson, 1932; Ghosh and Walocha, 2024; González, 2025). Of particular interest to the authors of this study is the Golgi-Cox method, which is an improved version of the original approach first developed by Camillo Golgi in 1873 (Dröscher, 1998; Zhong *et al*., 2019; González, 2025). The Golgi-Cox method, like its predecessor and other variations, has led to high structural visibility of neuronal structures to the level of dendritic spines as well as capturing changes in neuronal structure between control and disease models (Baloyannis, 2015; Kang *et al*., 2017). While many studies focus on cortical and hippocampal neurons (Flores *et al*., 2015; Zhong *et al*., 2019), how factors such as age and typically occurring daily variations in brain functions (known as circadian rhythms) may impact Golgi staining patterns across different brain regions is less studied. Regardless of the historical impact of the Golgi staining method, the reason why specific types of neurons are stained using this method remains unknown.

Here, we demonstrate that age, time of day, and timed environmental stimuli are important factors for the number of cells stained with the Golgi-Cox method. Specifically, we demonstrate that as neurons mature, more neurons are stained with the Golgi-Cox method, using 1-week-old, 2-week-old, and 2-month-old mice. Furthermore, in adult (2-month-old) mice, we demonstrate a circadian (time-of-day specific) pattern in the number of cells stained in hypothalamic nuclei involved in sleep and arousal. Finally, based on our cell number-related staining pattern and published data on neural activity in specific hypothalamic nuclei (SCN (Suprachiasmatic Nucleus) (Duy *et al*., 2020) and VLPO (Ventrolateral Preoptic nucleus) (Novak and Nunez, 1998; Miller *et al*., 2005)), we hypothesize that the Golgi-Cox staining pattern in adults is likely restricted to inactive neurons, which could be an explanation for both the sparse and variable staining pattern notoriously associated with the Golgi-Cox method. This study highlights the possibilities of using the Golgi-Cox method to answer a broad range of neuroscience questions related to development, circadian rhythms, and environmental changes.

## Methods

### Mice

We used CD-1 mice (*Mus musculus)* (from Charles River Laboratories), which are outbred mice with a reliable breeding consistency and litter size (Hart *et al*., 2018). Mice were housed and treated following NIH (National Institutes of Health), ARRIVE (Animal Research: Reporting of In Vivo Experiments) guidelines, and approved protocols by the OUACUC (Oakwood University Animal Care and Use Committee). Parental mice used for mating were three months old. For all experiments, we used 12:12 LD (light-dark), one of the most used light cycles to maintain and breed mice (Jennings *et al*., 1998). It consists of 12 hours of darkness and 12 hours of light (∼500 lux) that repeat every 24 hours. Food and water were provided *ad libitum,* and cage changes occurred every two weeks or more frequently as needed. Sibling littermate mouse pups were used for P7 and P14 time points (gender not determined) and adult (P60) males. Male mice were used in this study due to findings of increased variability in circadian-driven behaviors due to hormone variability associated with the 4-5 estrous cycle in female mice (Jud *et al*., 2005; Datta *et al*., 2016). Adult gender studies are being considered for the future.

### Experimental design

#### Histological analyses

Before tissue retrieval, mice were anesthetized using Avertin (20mg/mL) in phosphate-buffered saline (PBS) via intraperitoneal injection and a foot pad pinch to confirm sensitivity loss. The mice were then perfused (cardiac) with 1x PBS followed by 4% paraformaldehyde (PFA) and post-fixed in 4% PFA before performing Nissl or Golgi-Cox histochemistry. Brains were stored in the fridge (4°C) in DW (distilled water) (for the Golgi-Cox protocol) or 1x PBS (for the Nissl protocol) during the pre-staining period (this time varied from 1-14 days). Unless otherwise stated, reagents are from *Thermo Fisher, USA*.

#### Sample collection

Mice in each experimental group were co-housed and received consistent handling and environmental exposures for each experimental group as described in Figures 1-3. Mice in Figure 1 were sacrificed 2 hours following dark onset (ZT 14). ZT = *Zeitgeber (“time-giver”)* is a term used to describe the time of day relative to lights on (ZT 0) and lights off (ZT 12) in a 24-hour day-night cycle consisting of 12 hours of light and 12 hours of darkness (Duy *et al*., 2020). For Figure 2 experiments, mice were sacrificed at ZT 0, ZT 1.5, ZT 12, and ZT 13.5 as shown in Figure 2A. For Figure 3 experiments, after the 30-minute exposure to 500 lux light, mice were sacrificed starting at 45 minutes (with the anesthesia)-60 minutes (perfusion in motion) to achieve a 90-minute time period between the beginning of the light pulse and the cessation of conscious brain activity.

**Figure 1:**
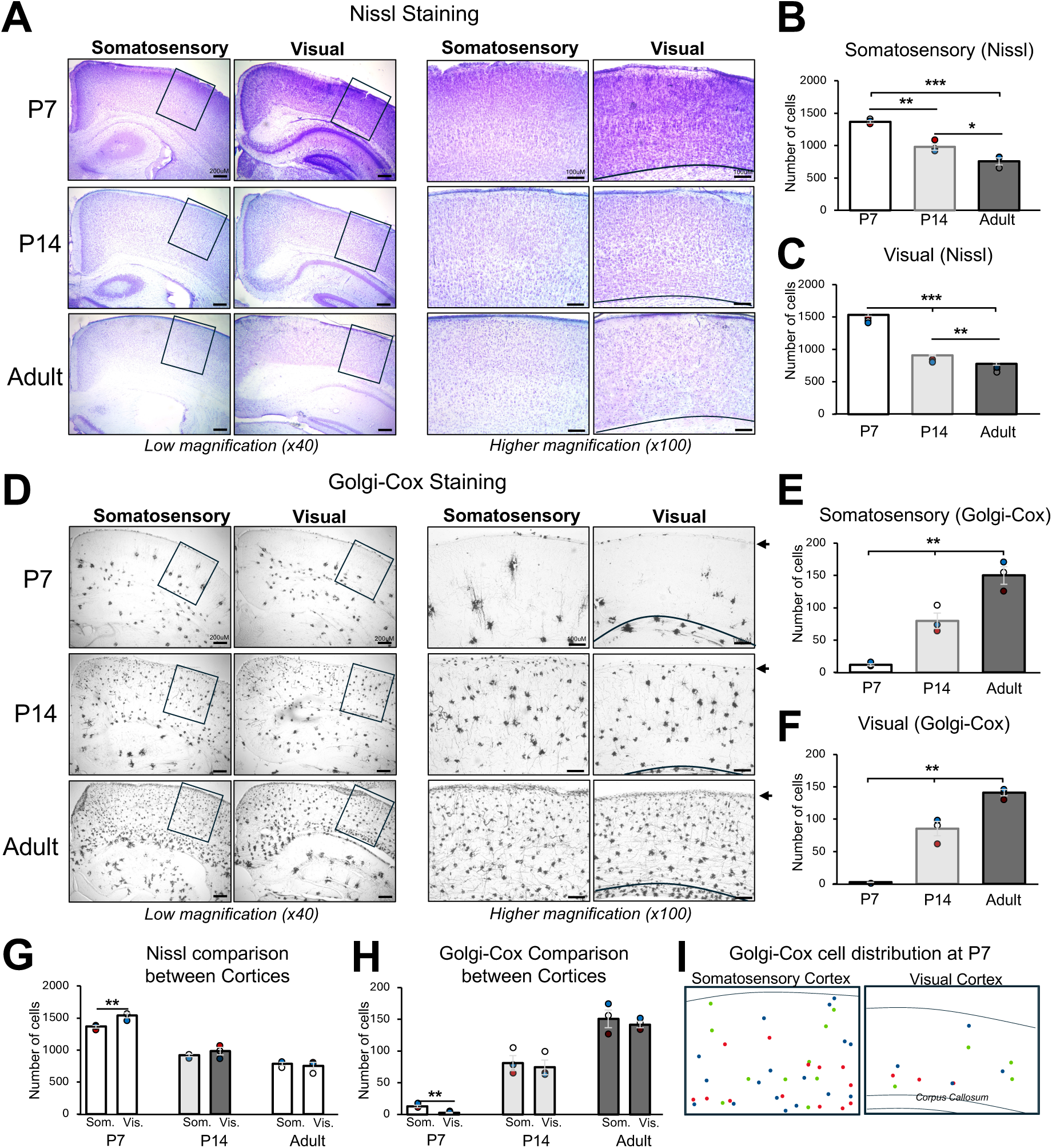
The number of Golgi-Cox-stained cells increases with age and correlates with functional differences in the somatosensory and visual cortices. A) Nissl staining of the somatosensory (primary) and visual (primary) cortices in P7, P14, and adult (P60) mice at low magnification (x40 -scale bars = 200 μM) with counting analysis area outlined with black boxes and at higher magnification (x100 -scale bars = 100 μM). The corpus callosum is separated from the visual cortex by a curved line towards the bottom of the higher magnification pictures. B) Cell count of Nissl-stained cells in P7, P14, and adult mice in the somatosensory cortex using the higher magnification images. *F*= 46.02, *P*= 2.3^-4^, average cell counts: 1365.14, 978.8, 756.4, and mean±s.e.m.: 21.4, 51.7, 55.2. C) Cell count of Nissl-stained cells at P7, P14, and Adult in the visual cortex using the higher magnification images. *F*= 461.98, *P*= 2.68^-7^, average cell counts: 1535.73, 909.69, 778.68, and mean±s.e.m.: 21.53, 12.04, 21.33. D) Golgi-Cox staining (grayscale) of the somatosensory and visual cortices in P7, P14, and adult mice at low magnification (x40 -scale bars = 200 μM) with counting analysis area outlined with black boxes and at higher magnification (x100 -scale bars = 100 μM). The corpus callosum is separated from the visual cortex by a curved line towards the bottom of the higher magnification pictures. Black arrows in the higher magnification visual cortex images indicate the location of the molecular layer. E) Cell count of Golgi-Cox-stained cells in P7, P14, and adult mice in the somatosensory cortex using the higher magnification images. *F*= 44.08, *P*= 2.58^-4^, average cell counts: 12, 80, 149.86, and mean±s.e.m.: 1.74, 12.04, 13.25. F) Cell count of Golgi-Cox-stained cells in P7, P14, and adult mice in the visual cortex using the higher magnification images. *F*= 101.54, *P*= 2.36^-5^, average cell counts: 2.83, 85.5, 140.92, and mean±s.e.m.: 2.83, 11, 4.67. G) Comparison between the number of Nissl-stained cells in the somatosensory and the visual cortices at P7, P14, and adult mice. *F*= 68.48, *P*= 2.11^-8^, average cell counts and Ranges: as stated in B, C, E, and F. H) Comparison between the number of Golgi- Cox-stained cells in the somatosensory and the visual cortices at P7, P14, and adult mice. *F*= 46.3, *P*= 1.96^-7^, average cell counts and Ranges: as stated in B, C, E, and F. I) Red, green, and blue dots in each image (somatosensory cortex and visual cortex) depict the location of Golgi-Cox-stained cells from three mice overlaid. For all experiments: *n*=3 mice, with each animal’s cell count average plotted with a white, red, or blue circle. Error bars are mean±s.e.m.; P-values are **P<0.01, ***P<0.001.

**Figure 2:**
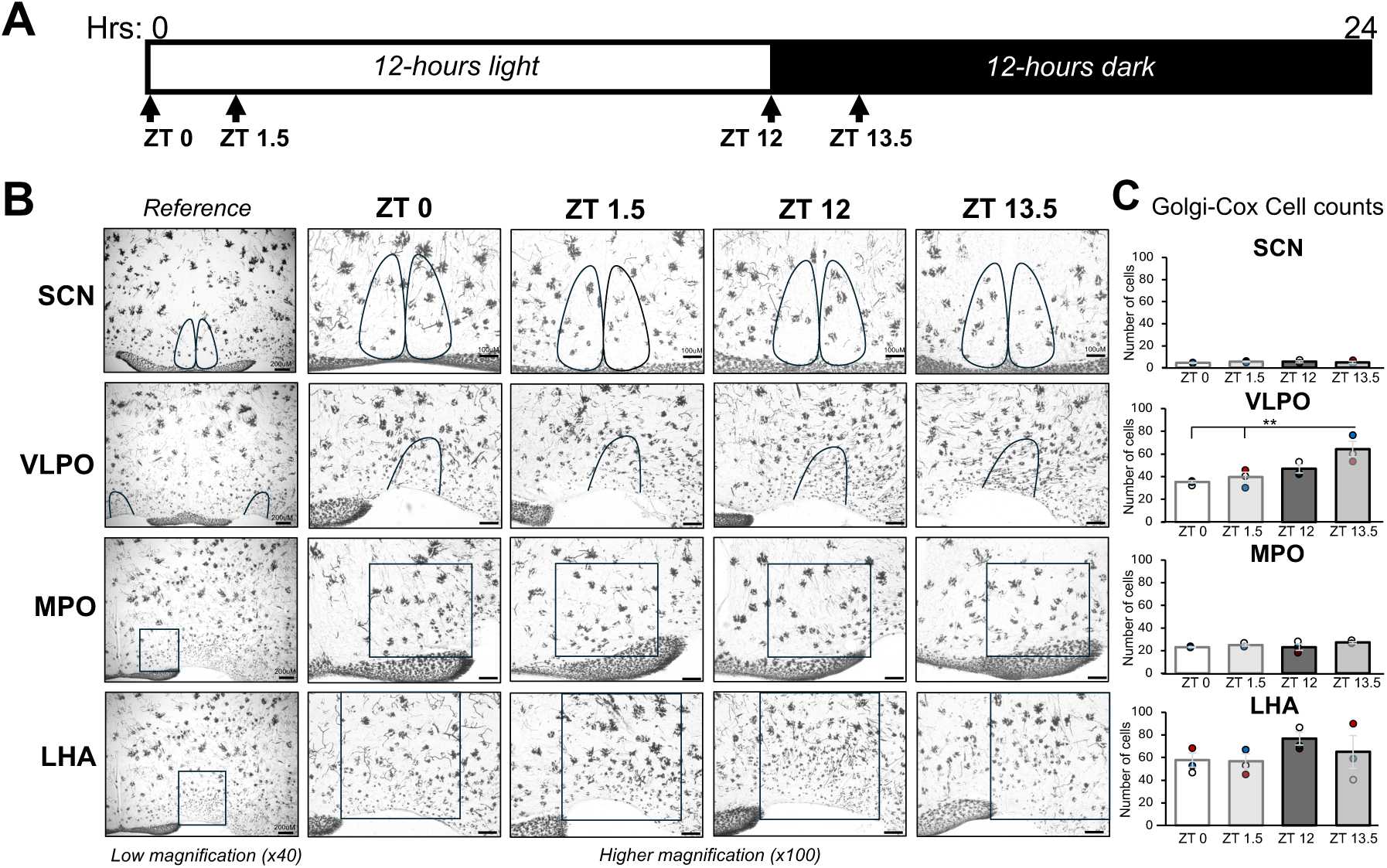
The number of Golgi-Cox-stained cells varies in a time-of-day dependent manner in some brain regions. A) Schematic of tissue collection timepoints during a 12-hr light: 12-hr dark cycle with the times during the day: ZT 0 = lights on, ZT 1.5 = 1.5 hrs after lights on; and times during the night: ZT 12 = lights off, ZT 13.5 = 1.5 hrs after lights off. B) Golgi-Cox staining (grayscale) in the SCN, VLPO, MPO, and LHA, at lower magnification (x40 -scale bars = 200 μM) and higher magnification at ZT 0, ZT 1.5, ZT 12, and ZT 13.5 (x100 -scale bars = 100 μM), with each region of focus outlined in black. C) Cell count of Golgi-Cox-stained cells in the SCN, VLPO, MPO, and LHA of the regions outlined in the higher magnified images. **SCN:** *F*= 0.57, *P*= 0.65, average cell counts: 4.44, 5.6, 5.5, 4.88, and mean±s.e.m.: 0.24, 0.4, 0.85, 1.06. **VLPO:** *F*= 7.94, *P*= 0.008, average cell counts: 35.43, 39.5, 46.94, 64.22, and mean±s.e.m.: 1.25, 4.54, 3.54, 6.83. **MPO:** *F*= 1.49, *P*= 0.28, average cell counts: 23.16, 25.16, 23.15, 27.22, and mean±s.e.m.: 0.33, 0.97, 2.87, 0.88. **LHA:** *F*= 1.05, *P*= 0.42, average cell counts: 57.8, 56.85, 782, 64.92, and mean±s.e.m.: 6.38, 6.43, 5.77, 14.45. For all experiments: *n*=3 mice, with each animal’s cell count average plotted with a white, red, or blue circle. Error bars are mean±s.e.m. P-value is **P<0.01.

**Figure 3:**
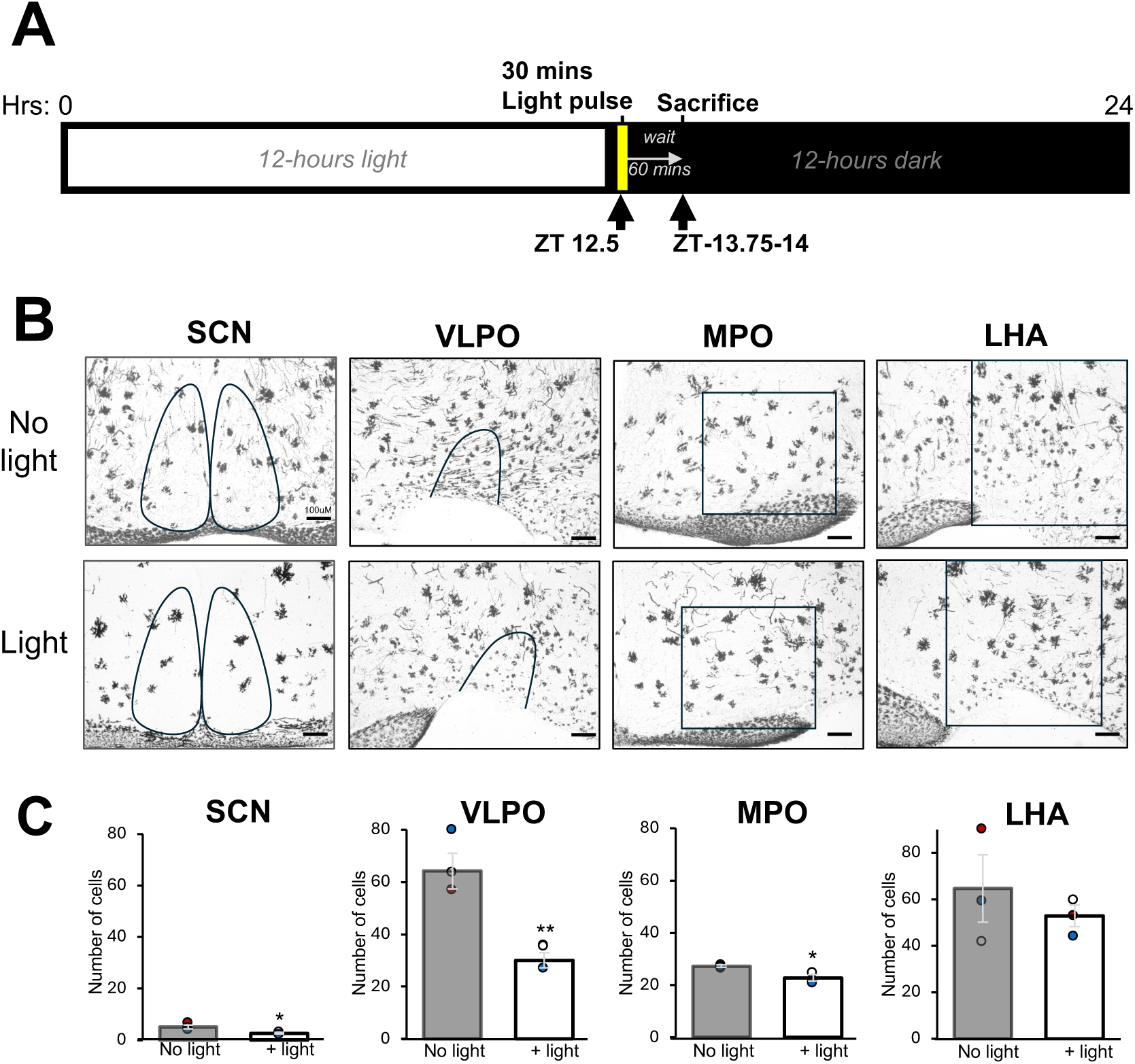
The number of Golgi-Cox-stained cells varies in response to light exposure at night in selected brain regions. A) Schematic of light pulse administration relative to the day-night cycle. B) Golgi-Cox staining (grayscale) in the SCN, VLPO, MPO, and LHA at higher magnification (x100 -scale bars = 100 μM) during the dark at ZT 13.5 (no light control) and ∼ZT13.5-14 following a 30-minute 500 lux light pulse at ZT 12.5 with each region of focus outlined in black. C) Cell count of Golgi-Cox-stained cells in the SCN, VLPO, MPO, and LHA of the regions outlined in the higher magnified images. **SCN:** *P*= 0.04, average cell counts: 5, 2.66, and mean±s.e.m.: 0.77, 0.33. **VLPO:** *P*= 0.01, average cell counts: 64.22, 30.12, and mean±s.e.m.: 6.83, 2.87. **MPO:** *P*= 0.02, average cell counts: 27.29, 22.77, and mean±s.e.m.: 0.39, 1.17. **LHA:** *P*= 0.48, average cell counts: 62.65, 52.83, and mean±s.e.m.: 14.49, 4.63. For all figures: *n*=3 mice, with each animal’s cell count average plotted with a white, red, or blue circle. Error bars are mean±s.e.m. P-value are *P<0.05, **P<0.01.

#### Nissl staining

The Nissl stain is used to stain neurons (although glial cells may be stained at times) in the brain (Einarson, 1932; Ghosh and Walocha, 2024). Brains being prepared for Nissl staining were sectioned (50 μM) using a vibratome (Lafayette instruments, USA) in 1x PBS at room temperature (22-24 °C). Slides with sectioned tissue were allowed to dry overnight at room temperature before staining. Slides were placed in a Coplin jar in 0.1% Cresyl Violet solution (Nissl stain) (*Abcam, UK*) for 5 minutes at room temperature. Slides were then dipped 4-5 times for 2 seconds each into differentiation solution (45 mL of 100% ethanol, 5 mL DW (distilled water), 5 drops of glacial acid), 100% ethanol, and Xylene. Slides were allowed to dry for 30 seconds before mounting (MM 24 mounting medium – *Leica, Germany*) and coverslipped.

#### Golgi-Cox staining

The Golgi-Cox stain is used to primarily stain neurons (Zhong *et al*., 2019) (although glial cells may be stained at times, depending on the protocol adjustments) (Narayanan *et al*., 2020). Our protocol is adapted from Zhong *et al*., 2019, in which neurons were the primary cell type visualized and modified for our study. The impregnation solution (solutions A (5% solution of Potassium dichromate (K_2_Cr_2_O_7_) in 100 ml DW), B (5% solution of Mercuric chloride (HgCl_2_) in 100 ml DW (heating required) and C (5% solution of Potassium chromate (K_2_CrO_4_) in 80 ml DW) and staining solutions (solutions E (20% ammonia solution) and F (1% sodium thiosulfate solution) were prepared and used as published in Zhong *et al*., 2019. Whole brains were placed in 24-well plate receptacles, covered with the impregnation solution, and kept in the dark at room temperature for 2 weeks. After rinsing the brains in DW 6 times (5 mins each time), the brains were kept in the fridge until sectioning (50 μM) using a vibratome. Slides were allowed to dry overnight before commencing the staining protocol, which ended with two dehydration steps in 95% then 100% ethanol, followed by Xylene (3 dips in each solution for 1-2 seconds). Slides were allowed to dry for 30 seconds before mounting (MM 24 mounting medium – *Leica, Germany*) and coverslipped.

### Data analysis methods

The Allen Brain Atlas was the primary source used to identify brain areas of interest for this study.

#### Locating primary somatosensory cortex

To consistently identify the primary somatosensory cortex, we first located the most rostral section that contained the hippocampus. We designated the middle of the primary somatosensory cortex to be diagonal from the outermost edge of the hippocampus (∼45-degree angle from the CA3 region) and captured images as shown in the outlined area in Figure 1A. This was done consecutively in the caudal direction for the next three 50 μM brain slices, totaling a 150 μM depth collected.

#### Locating primary visual cortex

To consistently identify the primary visual cortex, we first located the initial and prominent separation of the medial corpus callosum, which is found caudally in the brain. The primary visual cortex was designated as the vertical area above the middle apex of the corpus callosum (∼90° angle from the apex) and captured images as shown in the outlined area in Figure 1A. This was done consecutively in the caudal direction for three 50 μM brain slices, totaling a 150 μM depth collected.

#### Locating hypothalamic nuclei

Easily identifiable structural landmarks were used to reliably locate specific areas being studied as follows: <u>SCN</u>-the optic chiasm (shape and depression where the SCN is located) and the third ventricle, <u>MPO (medial</u> <u>preoptic area) and LHA (lateral hypothalamic area)</u> – these two regions are quite large, but we analyzed the region lateral to the middle of the SCN, and the <u>VLPO</u> – 150 μM rostral to the start of the SCN was the last section of the VLPO area analyzed. Three 50 μM brain slices were collected per area and condition, totaling a 150 μM depth collected.

#### Microscopy

We used a *Fisher Science Education* microscope, *AmScope (USA)* (MU1000-HS) camera, and the *AmScope* program to capture images at 4x (40 times magnified), 10x (100 times magnified), and 40x (400 times magnified) magnification for analysis. Scale bars are included in each representative image for size reference. Cortical pictures were captured with the molecular layer horizontally oriented (in contrast to the natural slanting seen in Figure 1A), while all other images were captured without adjusting the horizontal axis of the brain when viewing the cortex as the superior-most region and the hypothalamus as the ventral-most region.

#### Cell counts

Cell counts were manually assessed within the areas outlined in each figure. Nissl cells were identified and counted based on the presence of Nissl bodies (small purple dots) within a diffusely purple-stained cell body with a smooth and continuous cell membrane. Golgi-Cox-stained cells were identified and counted based on the classical, black-colored staining, along with confirming the presence of dendrites and axons. If neurons were found in an apparent cluster (indistinguishable cell bodies with numerous processes, which was more common in the P7 brains), the cluster was counted as one cell (this was very common throughout P7 brain tissue). The size of the area for each brain area studied was outlined before counting cells and was consistent between mice of the same age and across the experimental conditions.

#### Statistical analysis

Statistical analyses were conducted with Prism GraphPad, version 9.4.1. Three mice were used for each experiment; mice were not reused in different experiments. Data sets are based on three 50 μM sections with bilateral analysis (6 values per mouse-bilateral values collected for each 50 μM section x 3 mice). Where a comparison of three or more groups (such as Figures 1 and 2) was conducted, the one-way ANOVA analysis was performed with Tukey’s multiple comparisons post-test, because each data set was compared to the other two or three data sets. The two-sample, two-tailed T-test was used for Figure 3, where we compared two samples.

The alpha for all tests is 0.05. The distribution of data sets exhibited a Gaussian distribution of normality. The error bars represent the standard error of the mean. Descriptive statistics, including the range, *P*-values, mean±s.e.m., and *F*-values, are included in each figure legend.

#### Data availability

All data are available on reasonable request and directed to the corresponding author, Melissa E S Richardson.

## Results

### The age of mice influences the number of neurons stained using the Golgi-Cox method

During early development, more neurons are present in the brain, which are later pruned as neural circuits and are established with age and experience (Chechik *et al*., 1999; Sakai, 2020). The sense of touch, which is controlled by the somatosensory system, is more developed in 1-week-old mouse pups than vision, as mouse pups do not open their eyes until around 2 weeks of age, which allows us to examine functional differences and milestones during development (Larsen and Callaway, 2006; Shen and Colonnese, 2016). To determine the number of cells in the somatosensory (primary) and visual (primary) cortices during development, we used the Nissl stain to examine tissues retrieved from P7 mice pups, P14 mice pups, and adult (P60) mice (Figure 1A). A significant reduction in cell number was progressively observed with age from P7 to P14 to adult in both the somatosensory (Figure 1B; *F*= 46.02, *P*= 2.3^-4^) and visual cortices (Figure 1C; *F*= 461.98, *P*= 2.68^-7^). While the Nissl staining was useful for determining the number of cells at different developmental stages, the Golgi-Cox method is widely used to sparsely stain neurons in a manner that allows visualization of not only cell bodies but also axons and dendrites (Zhong *et al*., 2019). Although the use of Golgi staining during development is uncommon (Marin-Padilla, 2015), we were able to successfully use the Golgi-Cox staining method to visualize neurons in P7, P14, and adult mice (Figure 1D). A significant increase in cell number was progressively observed with age from P7 to P14 to adult in both the somatosensory (Figure 1E; *F*= 44.08, *P*= 2.58^-4^) and visual cortices (Figure 1F; *F*= 101.54, *P*= 2.36^-5^). A comparison between the somatosensory and visual cortex reveals significantly more Nissl-stained cells in the visual cortex at P7 and significantly fewer Golgi-Cox-stained cells in the visual cortex at P7 compared to the somatosensory cortex at P7 (Figure 1G; *F*= 68.48, *P*= 2.11^-8^ & 1H; *F*= 46.3, *P*= 1.96^-7^). Although the somatosensory and visual cortices are located in different regions of the brain and regulate different behaviors, there was no significant difference in cell counts for either the Nissl or Golgi-Cox staining in either P14 or adult mice (Figure 1G & H). There were two spatially relevant patterns observed during development with the Golgi-Cox staining: 1) the location of cells in the somatosensory cortex at P7 was more broadly distributed than that of the P7 visual cortex (Figure 1I) and 2) the molecular layer (indicated by the black arrows in Figure 1D) contains more cells with age in both the somatosensory and visual cortices. Together, these findings indicate that cell staining patterns for both the Nissl and Golgi-Cox methods are age-dependent and may be related to the functional maturity of the brain region being studied.

### The number of cells stained with the Golgi-Cox method varies in a time-of-day-specific manner in some brain areas

To further determine whether environmental conditions impacted the Golgi-Cox staining pattern, adult mice were sacrificed and histologically analyzed at four timepoints during the day-night cycle that were relevant to the sleep-wake cycle as follows: ZT 0 – potential time of sleep onset, ZT 1.5 – proteins related to sleep onset or neuronal activity related to sleep maintenance should be synthesized, ZT 12 – time of typical activity onset, ZT 13.5 – proteins related to activity onset or neuronal activity related to activity maintenance should be synthesized (Lee *et al*., 2018; Duy *et al*., 2020; Lara Aparicio *et al*., 2022) (Figure 2A). We selected hypothalamic nuclei known to be involved in circadian rhythms (SCN) (Duy *et al*., 2020), sleep (VLPO) (Sherin *et al*., 1996), and arousal (MPO and LHA) (McGinty and Szymusiak, 2001; Szymusiak and McGinty, 2008) (Figure 2B). No difference in the total number of Golgi-Cox-stained cells was found at different times of the day-night cycle for the SCN, MPO, or LHA (Figure 2C). However, a rhythmic pattern of Golgi-Cox-stained cells was detected in the VLPO with a significantly increasing cell count pattern from ZT 0 to ZT 13.5 (*F*= 7.94, *P*= 0.008). (Figure 2C). Together, these data indicate two key findings: 1) Golgi-Cox-stained cell number throughout the day can be consistent (indicating reproducibility of this method: SCN, MPO, and LHA), and 2) depending on the brain area, a circadian pattern of Golgi-Cox-stained cell number is detectable (VLPO).

### Light exposure during the early night decreases the number of Golgi-Cox-stained cells in light-responsive brain regions

Next, we wanted to determine whether Golgi-Cox-stained cell numbers would change in response to a stimulus at a specific time of the day. Previous studies demonstrate that light-responsive neurons, such as those found in the SCN, are activated in response to light exposure during the early night, as indicated by histological analysis of C-fos protein, an immediate early gene (Duy *et al*., 2020). Therefore, we hypothesized that if the Golgi-Cox staining is dependent on environmental events, even in a shorter window of time (90 minutes compared to 24 hours in figure 2), then we would detect a change in the number of Golgi-Cox-stained cells in the SCN and possibly other brain areas involved in sleep and arousal. To test this hypothesis, we administered a 30-minute light pulse at ZT 12.5 and waited 45-60 minutes, in a similar manner to how the C-fos studies were conducted, before sacrificing mice for histological analyses (Figure 3A). A significant decrease in the number of Golgi-Cox-stained cells was observed in the SCN (light responsive) (*P*= 0.04), VLPO (sleep induction) (*P*= 0.01), and MPO (arousal) (*P*= 0.02), but not LHA (arousal) (Figure 3B & C). These findings indicate that the number of neurons stained by the Golgi-Cox method relies on recent environmental changes, such as a response to light exposure at night, and that not all brain areas will respond to the stimulus in the same manner.

### The number of Golgi-Cox-stained cells is reciprocally correlated with published neural patterns of activity

Upon further examination of the data in Figures 2 and 3, we noticed additional patterns associated with the Golgi-Cox-stained cells. The SCN had a consistent number of cells across the four times of the day studied (Figure 2C). However, the location and distribution of the cells within the centrally located sub-regions, referred to as the core and shell, varied (Figure 4A and Lokshin *et al*., 2015). Specifically, we compared the middle section of the SCN at each time point by overlaying the cell position from each of the three mice and observed that the majority of the Golgi-Cox-stained cells were located in the shell region, at ZT 1.5 (*P*= 9.91^-4^) and ZT 12 (*P*= 0.01), which unveiled a previously unnoticed circadian pattern when the entire SCN was examined in Figure 2A-B (Figure 4A and 4B). Furthermore, mice that were exposed to the light pulse during the early night (Figure 3C) did not exhibit Golgi-Cox-stained cells in the core region (Figure 4A and 4B; *P*= 0.01).

**Figure 4:**
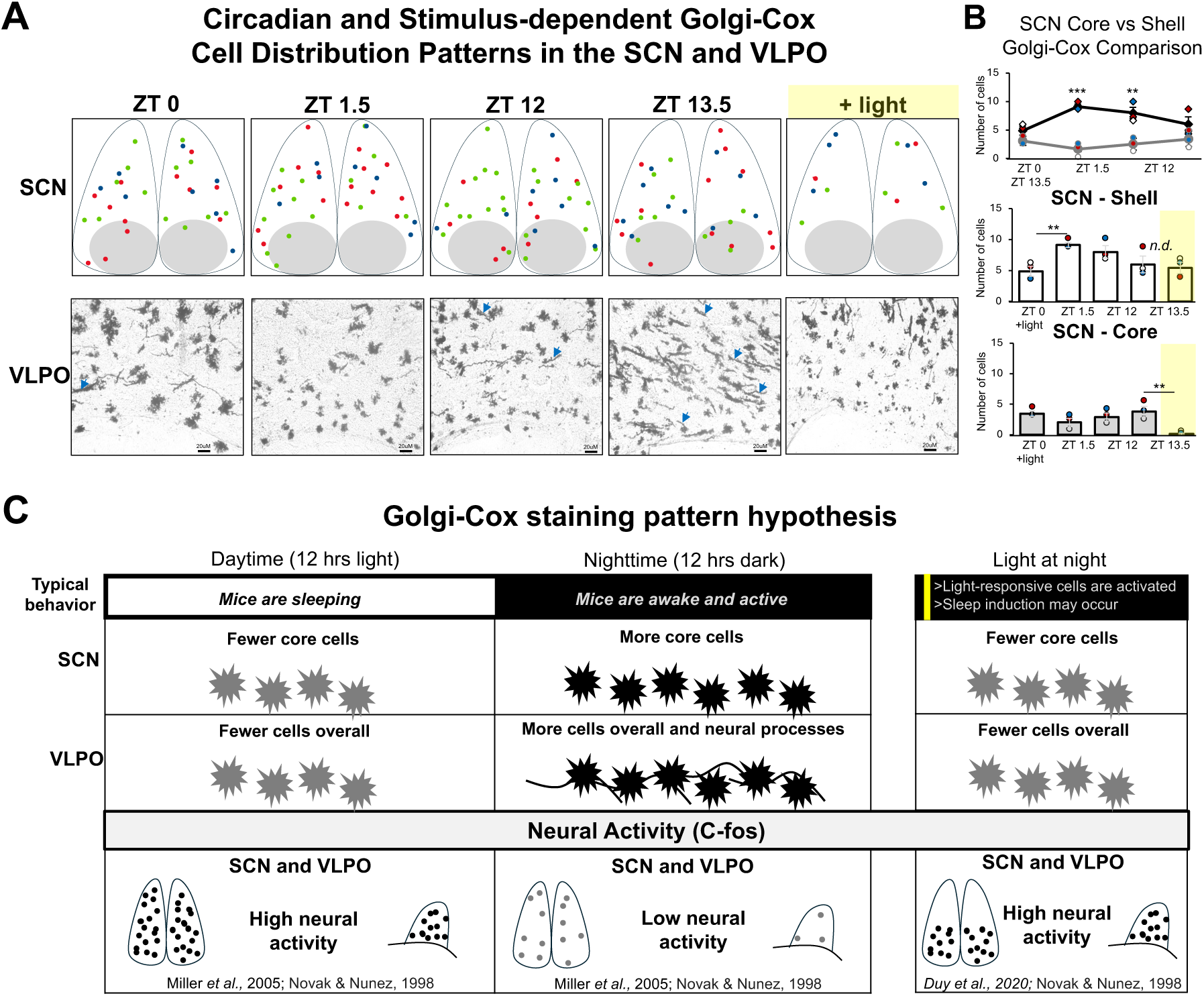
Fewer Golgi-Cox cells are stained at times of the day and conditions when neural responses are typically high. A) Location of Golgi-Cox-stained cells in the SCN and the presence of neural processes in Golgi-stained VLPO cells (grayscale) (blue arrows - x400 -scale bars = 20 μM) at ZT 0, 1.5, 12, 13.5, and with light at night (ZT 13.5-14). Red, green, and blue dots in each SCN image depict the location of Golgi-Cox-stained cells from three mice overlaid. The core region of the SCN is highlighted with gray circles. B) Average number of cells in the shell vs core regions of the SCN from the middle sections depicted in (A) at ZT 0 [*P*= 0.19, average cell counts: 4.88, 3.44, and mean±s.e.m.: 0.8. 0.44], ZT 1.5 [*P*= 9.91^-4^, average cell counts: 9.11, 2, and mean±s.e.m.: 0.44, 0.69], ZT 12 [*P*= 0.01, average cell counts: 8, 2.88, and mean±s.e.m.: 1.02, 0.67], ZT 13.5 [*P*= 0.23, average cell counts: 6, 3.77, and mean±s.e.m.: 1.34, 0.86], SCN shell (ZT 13.5) compared light at night (ZT 13.5-14), [*P*= 0.75, average cell count with light: 5.44, and mean±s.e.m.: 0.9], and SCN core (ZT 13.5) compared to light at night (ZT 13.5-14) [*P*= 0.01, average cell count with light: 0.11, and mean±s.e.m.: 0.11]. C) Comparison of Golgi-Cox-stained cell number and neural activity patterns in the SCN and VLPO at ZT 0, 1.5, 12, 13.5, and with light at night (ZT 13.5-14). Black and gray dots represent the comparative magnitude of neural activity in the SCN and VLPO, but they are not an actual representation. For all figures: *n*=3 mice, with each animal’s cell count average plotted with a white, red, or blue circle. Error bars are mean±s.e.m. P-value is *P<0.05, **P<0.01 and ***P<0.001.

The VLPO exhibited an increasing number of Golgi-Cox-stained cells from ZT 0 to ZT 13.5 (Figure 2C). We also noticed that the cell morphology differed between the day (ZT 0, ZT 1.5) and the night (ZT 12, ZT 13.5) time points, with more neural processes apparent on the cells at ZT 12 and ZT 13.5 (Figure 4A; blue arrows).

The impact of light exposure on the number of Golgi-Cox-stained cells (reduction) is opposite to published patterns of neural activity (increased, using C-fos, a neuronal activation marker) in the SCN (Duy *et al*., 2020) and VLPO (Novak and Nunez, 1998; Miller *et al*., 2005). In the SCN in particular, C-fos cell number increased dramatically in the core region of the SCN, while we observed a near-zero cell number of Golgi-Cox-stained cells (*P*= 0.01) using a similar experimental design (Duy *et al*., 2020). These observations led us to the hypothesis that the Golgi-Cox-stained cells in our studies are likely inactive (Figure 4C).

## Discussion

This study provides a multifaceted approach to understanding that the pattern of Golgi-Cox staining is not random and depends on the state of the brain relative to the environment. Furthermore, we provide a strong data set with low variability (as noted by the mean±s.e.m. values in the figure legends) that indicates consistent results within each experimental group. We acknowledge that our findings are limited to the brain areas we selected, but emphasize that even with such a selective sampling, our findings are impactful in their interpretation and potential applications.

Our comparison of two cortical regions (the primary somatosensory and visual cortices) with easily observable behaviors (touch and vision) allowed us to discover a noticeable developmental delay in the Golgi-Cox staining pattern during week 1 (P7) when eyes are still closed, but touch is active in mouse pups (Figure 1). Our data showing a reduction in the Nissl-stained cell number with increased age is consistent with previous studies that show mass pruning of neurons during the first two weeks of development (Zeiss, 2021). The similarity in the number of Golgi-Cox-stained cells in both the somatosensory (Larsen and Callaway, 2006) and visual cortices (Shen and Colonnese, 2016) at P14 may indicate that a similar maturity level has been achieved. Though these are two different cortices, research indicates increased neural activity in the primary visual cortex just before the eyes open (Shen and Colonnese, 2016). We also noticed that neurons in P7 mice are structurally distinctive from neurons in P14 and adult mice, which requires further investigation in a future study. The molecular layer, which is involved in integrative functions within the cortex, also shows an age-specific staining pattern, with more cells visible with increased age (Genescu and Garel, 2021). There are many other cortical areas and brain regions to be examined in the future to determine the pervasiveness of this phenotype.

In our adult-only studies, we focused on sleep and arousal, relative to the 24-hr day-night cycle, due to the relevance of circadian-driven behaviors and the authors’ familiarity with brain areas regulating these typically occurring functions. While the SCN and MPO did not show a circadian variation in cell number and had notably fewer cells overall at each time point (Figure 2C), upon further analysis, the SCN did show circadian variation in subregions (Figure 4A & B). The comparably small number of cells stained in the SCN and MPO may be too few to show any potential circadian variation and indicate the range of cell density by specific region (some areas, such as the VLPO and LHA, have more cells stained than others) (Figure 2). While there are many more brain regions to be examined that are influenced by circadian rhythms, this data indicates that the time of day at which tissue for the Golgi-Cox staining is retrieved is relevant for the number of cells that may be observed.

Light at night is known to change sleep and arousal behavior and activate cells in brain areas such as the SCN (Duy *et al*., 2020), VLPO (Novak and Nunez, 1998; Miller *et al*., 2005), and the POA (preoptic area) (Zhang *et al*., 2021), and is correlated with a reduction in wheel-running activity and sleep induction (Hubbard *et al*., 2013; Liu *et al*., 2024). We show that light exposure during early night reduced cell numbers in the SCN, VLPO, and MPO but not the LHA (Figure 3). We were intrigued to find in our studies that, in using a similar experimental design to previous light-induced C-fos IHC studies, there were fewer cells in the SCN, VLPO, and MPO (Figure 3). Upon further examination of the SCN, we discovered that a reduction in cells occurred in the core region of the SCN, an area previously characterized to exhibit dense neural response to light at night (Figure 4 and Duy et al., 2020). This data, along with our findings that the shell SCN cells do exhibit a circadian rhythm, bolsters our conclusion that although the total number of cells stained in the SCN was consistent in Figure 2 (time-of-day study), the sub-location of the cells and the response to light yielded significant and physiologically relevant data (Figure 4A & B). Furthermore, in the VLPO, not only were fewer cells stained following light exposure at night, but the distinctive neural processes observed at ZT 13.5 were absent (Figure 3B and Figure 4A), which demonstrates that the Golgi-Cox stain captured structural neural changes in response to the light exposure at night. We did not notice any additional phenotypes in the region of the MPO we examined, which is part of a relatively smaller fraction of the larger POA (compared to the SCN and the VLPO, which are smaller structures). The LHA, though light responsive, regulates many functions, including arousal, but did not show a significant decrease in Golgi-Cox-stained cells in response to light at night, and like the POA, is a relatively large structure that has functional subregions (Gazea *et al*., 2021).

Together, these data, especially the findings in the SCN and VLPO, have led us to believe that in adults, the Golgi-Cox staining captures the structure of quiescent neurons for the following reasons: 1) the VLPO, which has fewer Golgi-Cox-stained cells during the day when mice are sleeping (vs night when mice are awake) is known to have more neural activity during the day, 2) the SCN (Duy *et al.,* 2020), VLPO (Novak and Nunez, 1998; Miller *et al*., 2005), and MPO (Zhang *et al*., 2021) exhibit fewer Golgi-Cox-stained cells following light exposure at night, but are known to have more neural activity following light exposure at night in published work, (Figures 3 and 4).

Our study is not without limitations. For instance, although the gender of the P7 and P14 mouse pups was not determined, studies show that glial and newly born cells in the brain are not gender-dependent or gender-ambiguous during the first 2 weeks of mouse pup development (Weinhard *et al*, 2018; Yagi *et al*., 2020). Furthermore, in our study, we used only male adult mice and 3 mice per experimental condition (with 6 technical replicates per mouse) to minimize the number of mice sacrificed to address our research questions and reduce potential gender-based variability in our adult brain studies, since we wanted to establish a baseline phenotype using the Golgi-Cox method. Using female mice in the future will provide valuable information, and we will work to develop the necessary tools needed to appropriately conduct adult gender comparison studies. Thus, we acknowledge that our study requires future expansion to make more comprehensive conclusions based on gender and age.

## Conclusion

This study is an important step toward understanding how the historical Golgi (-Cox) method of visualizing neurons works and redirecting its use to answer more targeted research questions cost-effectively. While there are many more brain regions and environmental conditions to explore using the Golgi-Cox method to enhance our understanding of the mechanism behind its staining action, this study provides much progress towards this goal. Future studies that focus on studying the entirety of brain areas or examining neural structure at different times of the day (as this method is classically used) will deepen our understanding of how the brain works using this method. Other variations to the Golgi method may yield different results and have not been explored yet. If it is proven in fact that only quiescent cells are stained by the Golgi-Cox method, use of this method in conjunction with other methods that identify neural activity could provide insight into the complementarity of how the brain uses excitatory and inhibitory responses to regulate a range of behaviors.

Importantly, the Golgi method will continue to be an excellent tool to capture the location and high-resolution structure of more functionally relevant neural data.

## Acknowledgments

Special thanks to Juliet Martin for support with reagent preparation, Shiraz Davis for leading mouse care, and the Department of Biological Sciences at Oakwood University for supporting this research.

## Author contributions

**Melissa E S Richardson**: Conceptualization, Methodology, Formal analysis, Investigation, Writing-all drafts, Visualization. **Lauren Barnwell:** Investigation, Writing-Original draft.

